# When perfect is a warning: explanations expose shortcuts that accuracy hides in cancer gene-expression classifiers

**DOI:** 10.64898/2026.11.02.763649

**Authors:** Hamail Ayaz, Pooja Verma, Muhammad Mahmood Ali, Oladosu Oyebisi Oladimeji, Ayodeji Olusegun Ibitoye

## Abstract

When the classes in a training set come from different data sources, a classifier can score near-perfectly while learning where a sample came from. We show this for cancer classifiers trained on tumours from The Cancer Genome Atlas and normal tissue from the Genotype-Tissue Expression project. Six model families, including a tabular foundation model, reached an area under the curve of 1.000 yet called 80–97% of tumour-adjacent normal tissue cancer, and 100% in an independent cohort; a few matched normal samples per tissue removed most of this shortcut. Near-perfect accuracy could not tell a shortcut from an honestly easy task, flagging 87–92% of models trained on such tasks. We introduce a label-free score, computed from model explanations, that uses the fact that a source difference looks the same in every tissue whereas cancer changes do not. It raised no false alarms, ranked models by their real reliance on the source, transferred to unseen model types and data, and failed where its theory predicts.

## Introduction

Public molecular data are now large enough to train machine learning models for cancer diagnosis without collecting a single new sample. The two largest bulk RNA-sequencing resources are The Cancer Genome Atlas (TCGA), which holds tumours and a limited number of tumour-adjacent normal tissues^1^, and the Genotype-Tissue Expression (GTEx) project, which holds normal tissues from post-mortem donors^2^. A classifier that has learned cancer should rely on changes that tumours share, such as faster cell division, invasion of the surrounding tissue and loss of normal tissue function^3^. No single gene marks every tumour at the level of RNA^4^. Whether a classifier has learned cancer therefore cannot be read from one marker; it has to be judged from all the genes it uses. TCGA and GTEx supplies what each lacks therefore, they are routinely combined, and uniform reprocessing of both. The same pipeline has made this combination easy^5,6^. Classifiers trained on tumours from one resource and normal tissue from the other report accuracies of 97–99%^7,8^, and models trained only on GTEx have been used to flag TCGA tumours as anomalies^9^.

In this design, however, the label and the data source are the same variable: every tumour comes from one source and almost every normal sample from the other. Differences between sources are a well-known problem in high-throughput biology^10^, and when they line up with the label they inflate performance estimates^11^. TCGA and GTEx differ in donors, sample collection (surgical resection versus post-mortem donation), handling and sequencing centres, and tumour-adjacent tissue is itself not the same as healthy tissue^12^. Uniform reprocessing removes differences in the computational pipeline, but not these. A classifier can therefore reach near-perfect accuracy by recognising where a sample came from rather than whether it is cancer. Such “Clever Hans” behaviour, in which a model is right for the wrong reason^13,14^, is well documented in medical imaging, where models have learned hospital markers, scanner signatures and tissue-source sites instead of disease^15–17^, and it has recently been shown in unsupervised models^18^.

Detecting such a shortcut is harder than it sounds. A held-out test set drawn from the same two sources contains the same confound, so accuracy stays near-perfect. Existing detection methods need information that is often missing: labels for the suspected shortcut attribute^19^, external data or targeted retraining^20^, or a pool of unlabelled deployment samples that already contains the failing cases^21^. Explanation methods can reveal Clever Hans behaviour^13,18,22^, but they are usually read by eye, and post-hoc explanations can miss spurious signals that the user does not already suspect^23^. In practice, researchers fall back on an informal rule: an AUC that looks “too good to be true” is suspicious. This rule has never been tested, and it has an obvious weakness: some tasks are simply easy.

Here we show that near-perfect accuracy cannot separate a shortcut from an easy task, and that a label-free score computed from model explanations can (Fig. 1). We use cancer-versus-normal classification from TCGA tumours and GTEx normals as a case study, and replicate the key results on a second, independently processed cohort, the Clinical Proteomic Tumor Analysis Consortium (CPTAC)^24^. We make four main contributions:

**Figure 1:**
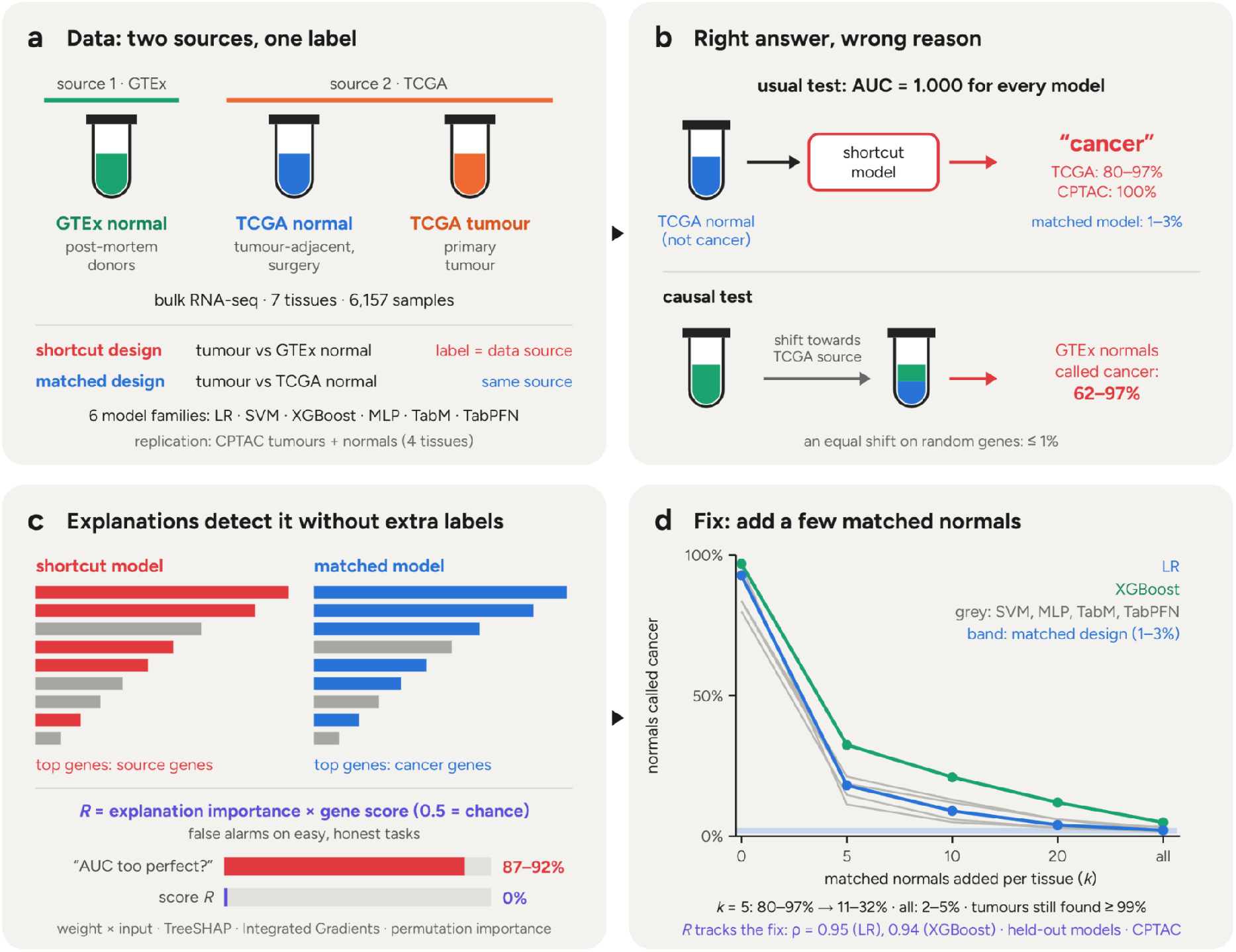
Explanations expose a data-source shortcut that near-perfect accuracy hides. **a**, TCGA supplies tumours and tumour-adjacent normal tissue; GTEx supplies normal tissue from post-mortem donors. In the shortcut design, tumours from TCGA are compared with normals from GTEx, so the label and the data source coincide; in the matched design, both classes come from TCGA. Six model families were trained on each design, and CPTAC was used for replication. **b**, On the usual test every shortcut model is near perfect, yet it calls most tumour-adjacent normals cancer (TCGA 80–97%, CPTAC 100%; matched models 1–3%). Shifting GTEx normals towards the TCGA source on the source genes makes 62–97% of them “cancer”; the same shift on random genes changes at most about 1%. **c**, Explanations show which genes a model uses. The source-reliance score *R* weights each gene’s importance by how source-like it is (0.5 = chance). On easy, honest tasks the “AUC too perfect” rule raises false alarms for 87–92% of models; *R* raises none. **d**, Adding a few matched normals per tissue to training removes the shortcut; *R* follows this change (Spearman *ρ* = 0.95 for logistic regression, 0.94 for XGBoost). Band, matched design. Panels a–c are schematic (explanation bars are illustrative); the numbers in b–d and the curves in d are from the data (means over 5 seeds; Figs. 2–4).

1. **First**, classifiers from six model families, from logistic regression to a tabular foundation model^25^, reach an AUC of 1.000 yet call 80–97% of tumour-adjacent normal tissue “cancer”; on CPTAC the rate is 100%.
2. **Second**, explanations show that these models rely on genes that separate the two data sources rather than cancer from normal tissue, whereas models trained on matched tumour and normal tissue from one source rely on cancer genes.
3. **Third**, adding a handful of matched normal samples per tissue to training removes most of the shortcut while keeping tumour detection intact, and works better than correcting the data with the same samples.
4. **Fourth**, we introduce a label-free shortcut score that needs no matched samples and no deployment data. It rests on a simple observation: a data-source difference tends to look the same in every tissue, whereas cancer changes are largely tissue-specific. On models trained on honestly easy tasks, the “too good to be true” rule raised false alarms for 87–92% of models, whereas our score raised none; the score also ranked models by how strongly they relied on the shortcut, generalised to model types it was never developed on, and replicated on CPTAC for linear models. We also report where it fails, and why.

The problem is not specific to cancer. Whenever cases and controls come from different studies, hospitals, biobanks or assays, a model can learn the origin of a sample instead of its biology. Our results suggest that accuracy alone should not be read as evidence that a model has learned the intended signal, and that explanations, used quantitatively rather than by eye, can provide that evidence without extra labels.

## Results

Following the experimental design and validation framework established above, we evaluated whether high-performing molecular classifiers learn true biological signals or data-source arti-facts. We conducted a systematic empirical analysis across six diverse model families, multiple tissue strata, and independent validation cohorts to demonstrate the ubiquity of source-based shortcut learning, validate remediation strategies, and test our explanation-based score.

### Accuracy and the data-source shortcut

We trained six model families, from logistic regression to the tabular foundation model TabPFN, to separate TCGA tumours from GTEx normal tissue of the same organ (the shortcut design; Fig. 1a). On the usual held-out test, every model was close to perfect: the AUC between tumours and GTEx normals was 1.000 for all six families (95% CI 1.00–1.00), and at least 99.9% of tumours were called cancer (Fig. 2a and Table 1).

**Table 1:** The shortcut and its fix across model families and two datasets. AUC: tumour vs GTEx normal (the usual test) and tumour vs tumour-adjacent normal, for the shortcut design. Tumour-adjacent normals called cancer: by the shortcut design; by the same model with a new decision threshold set with 5 matched normals per tissue; after retraining with *k* = 5 or all matched normals per tissue; and by the matched design. Tumours missed: share of tumours lost by the new threshold (retrained models keep tumour detection at *≥* 99%). GTEx normals called cancer: by the matched design, and by the mixed design (shortcut design plus all matched normals). Means over three seeds. Dashes: not computed (no new threshold was set on CPTAC).

|  |  | AUC (shortcut design) |  | Tumour-adjacent normals called cancer (%) |  |  |  |  | Tumours missed (%) | GTEx normals called cancer (%) |  |
|---|---|---|---|---|---|---|---|---|---|---|---|
|  |  |  |  |  | New | Retrained |  |  |  |  |  |
| Data | Model | Usual | Adjacent | Shortcut | threshold | *k* = 5 | All | Matched | New threshold | Matched | Mixed |
| TCGA | LR | 1.00 | 0.98 | 93 | 10 | 18 | 2 | 2 | 9 | 29 | 0 |
|  | SVM | 1.00 | 0.98 | 84 | 8 | 19 | 2 | 3 | 6 | 36 | 0 |
|  | XGBoost | 1.00 | 0.93 | 97 | 15 | 32 | 5 | 3 | 17 | 17 | 0 |
|  | MLP | 1.00 | 0.93 | 80 | 27 | 15 | 4 | 1 | 3 | 9 | 0 |
|  | TabM | 1.00 | 0.97 | 84 | 13 | 21 | 3 | 1 | 7 | 9 | 0 |
|  | TabPFN | 1.00 | 0.96 | 96 | 11 | 11 | 2 | 1 | 14 | 5 | 0 |
| CPTAC | LR | 1.00 | 0.89 | 100 | – | 3 | 1 | 1 | – | 65 | 0 |
|  | XGBoost | 1.00 | 0.48 | 100 | – | 24 | 1 | 1 | – | 90 | 0 |

**Figure 2:**
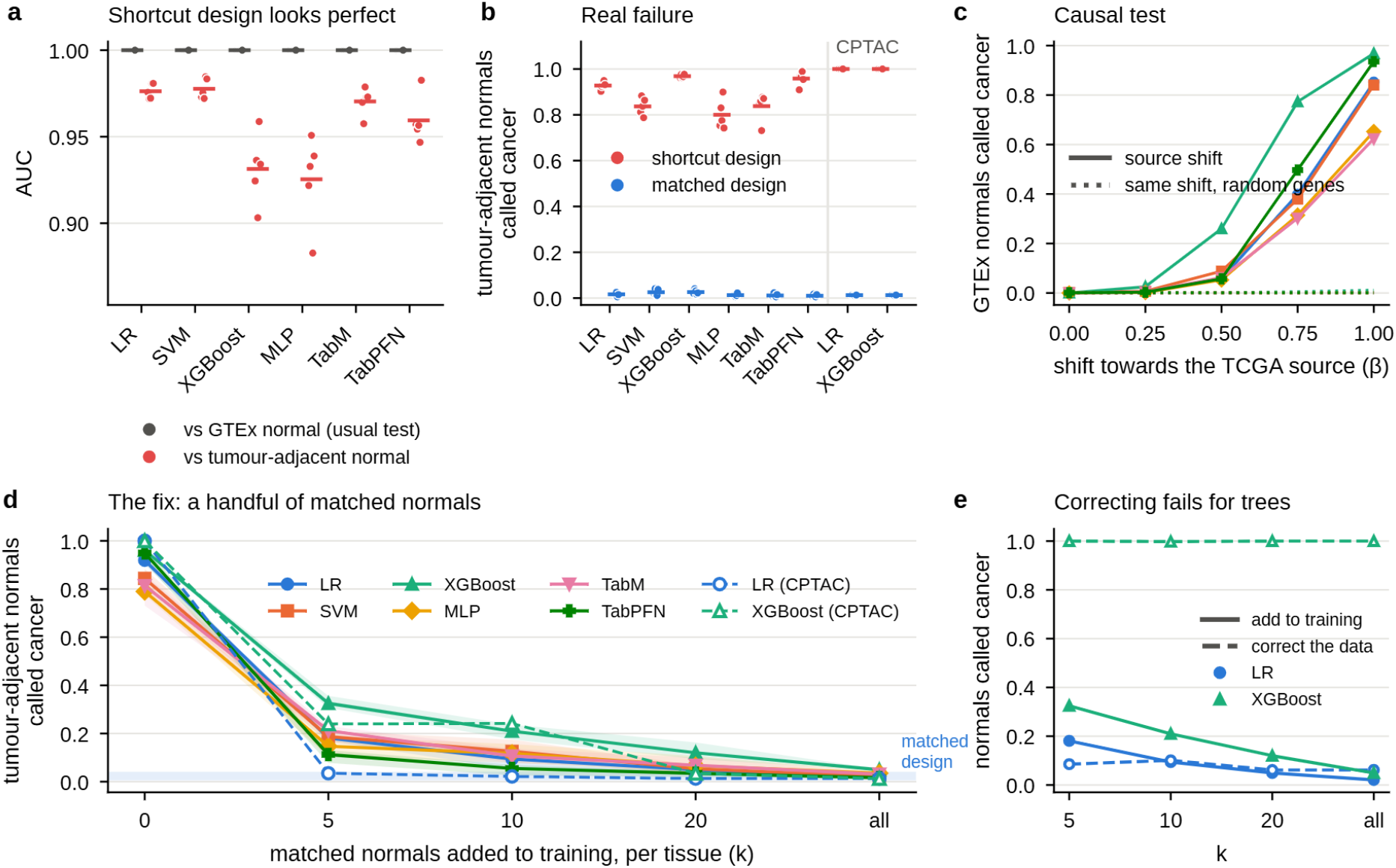
Near-perfect classifiers learn the data source, and a few matched normals remove the shortcut. **a**, AUC of shortcut-design models (tumour vs GTEx normal) on the usual test (grey) and against TCGA tumour-adjacent normals (red); dots are seeds, bars are means. The y axis starts at 0.85. **b**, Share of tumour-adjacent normals called cancer by the shortcut design (red) and the matched design (blue), on TCGA (six families) and CPTAC (logistic regression and XGBoost). **c**, Causal test: share of held-out GTEx normals called cancer after shifting them by a fraction *β* of the source difference towards TCGA (solid), or by the same shift with genes permuted (dotted). **d**, Share of tumour-adjacent normals called cancer after adding *k* matched normals per tissue to training, on TCGA (filled, solid; shading: range over seeds) and CPTAC (open, dashed). Blue band: matched design. **e**, Adding the matched normals to training (solid) versus using them to correct the data (dashed), for logistic regression and XGBoost. Colours denote model families throughout.

The same models failed on tumour-adjacent normal tissue from TCGA. They called 80% (MLP; 95% CI 73–85%) to 97% (XGBoost; 93–99%) of these normals ‘cancer’ (Fig. 2b). Models of the same families trained on tumours and tumour-adjacent normals from TCGA (the matched design) called only 1–3% of the same normals cancer (P < 0.001 for every family, Fisher’s exact test). The failure was hidden by the usual test, yet visible in it once the right comparison was made: against tumour-adjacent normals, the AUC of the shortcut models fell to 0.93– 0.98. On a second, independently processed cohort (CPTAC), the failure was complete: logistic regression and XGBoost called 100% of CPTAC tumour-adjacent normals cancer in every seed, and the AUC of XGBoost between CPTAC tumours and normals was 0.48, no better than chance (Table 1). Models of the matched design called 1% of them cancer.

Two explanations fit this result: the models learned the data source, or tumour-adjacent tissue is itself partly abnormal^12^. To tell them apart, we moved held-out GTEx normals towards the TCGA source, without adding any cancer signal (Fig. 2c). At a full shift, the shortcut models called 62% (TabM; 95% CI 57–67%) to 97% (XGBoost; 94–98%) of these GTEx normals cancer. The same shift with its genes randomly permuted, which keeps its size but breaks its direction, changed almost nothing (0.0–1.1%; P < 0.001 for every family). The data source alone is therefore enough to make these models call normal tissue cancer. This was not a consequence of how genes were selected: the failure remained when models were trained on 2,000 random expressed genes or the 2,000 most highly expressed genes (Supplementary Table B).

### Matched samples and the shortcut

If the shortcut arises because the label and the source coincide, breaking the coincidence should remove it. We added *k* TCGA normals per tissue to the training set of the shortcut design (Fig. 2d). Five per tissue (35 samples in total) cut the share of tumour-adjacent normals called cancer from 80–97% to 11% (TabPFN; 95% CI 7–17%) to 32% (XGBoost; 26–40%), and adding all available matched normals brought it to 2–5%, the level of the matched design. Tumour detection was kept throughout (99.1–99.7% of tumours called cancer at *k* = 5). The fix replicated on CPTAC, where five matched normals per tissue reduced the failure of logistic regression from 100% to 3%, and twenty reduced that of XGBoost to 3% (Fig. 2d). Training with both sources also avoided a weakness of the matched design: models trained only on TCGA called 5–36% of GTEx normals cancer (65–90% on CPTAC), whereas the mixed design called none (Table 1).

The same matched samples could instead be used without retraining. Choosing a new decision threshold with them lowered the share of normals called cancer to 8–27%, but at the cost of missing 3.0–16.8% of tumours, against 0.3–0.9% after retraining (Extended Data Fig. 2). Using them to correct the data rather than to train the model worked for logistic regression, and at *k* = 5 even better than retraining (8.5% vs 18.1% of normals called cancer; P = 0.021), but failed completely for XGBoost, which still called 100% of normals cancer at every *k* (Fig. 2e). A tree ensemble can split on a gene at a value that a mean shift does not move, so correcting the data does not remove what the model has learned. Adding matched samples to training was the only approach that worked for every family.

Each step of this series also gives a model with a known, graded reliance on the source, from complete (*k* = 0) to almost none (all). We use this ladder below to test whether a shortcut can be measured without matched samples.

### Explanations of the shortcut

Matched samples fix the shortcut; however, most studies that combine sources do not have them, and without them a shortcut cannot be seen in accuracy. We therefore asked whether the models’ explanations reveal what they use. For each gene, we measured its source effect (TCGA normal vs GTEx normal) and its cancer effect (TCGA tumour vs TCGA normal), and compared them with the genes each model relied on most.

The two designs used different genes (Fig. 3a,b). Of the 50 genes that shortcut logistic regression relied on the most, 36 had a larger source effect than cancer effect (median |*d*| 1.6 vs 0.8); of the 50 used most by the matched model, only 15 did (median 0.7 vs 1.0). The same pattern held for every model type and seed (Fig. 3c). Each shortcut design model relied more on source genes than any matched design model of the same type, and the matched-design models shifted their reliance towards cancer genes, most clearly for XGBoost and TabM.

**Figure 3:**
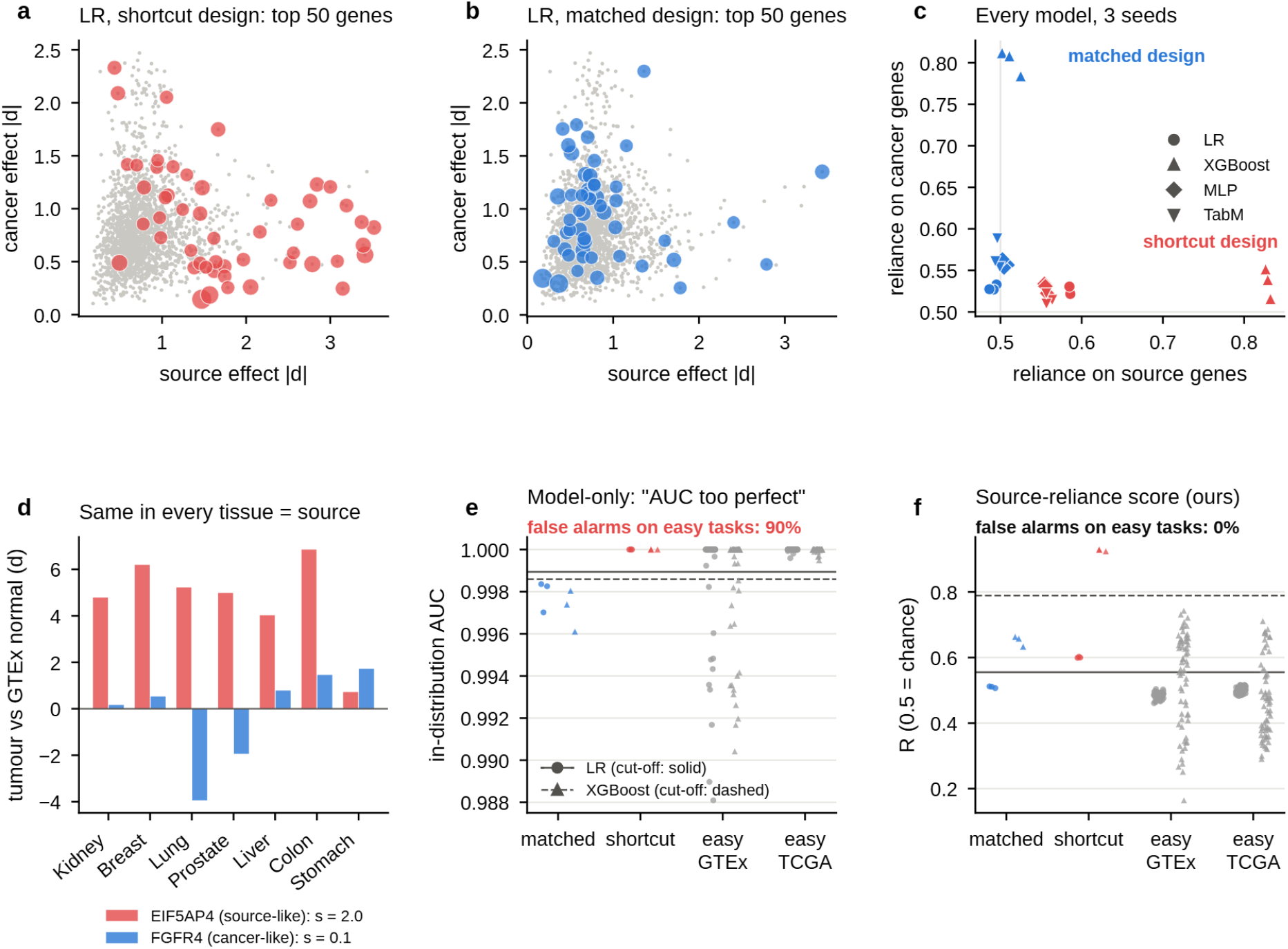
Explanations reveal the shortcut, and a label-free score turns this into a check that accuracy cannot provide. **a, b**, Source effect and cancer effect (|*d*|, mean over tissues) of every gene (grey); the 50 genes that logistic regression relies on most are coloured, with size proportional to importance, for the shortcut (a) and matched (b) designs. **c**, Importance-weighted reliance on source genes and on cancer genes for every model and seed; colour: design; shape: model type. **d**, The idea behind the score: difference between tumours and GTEx normals in each tissue for a source-like gene (*EIF5AP4*) and a cancer-like gene (*FGFR4*); *s*, gene score (Eq. 2). **e, f**, In-distribution AUC (e) and source-reliance score *R* (f) for matched-design and shortcut-design models and for negative controls (easy tasks with no source difference: tissue vs tissue within GTEx or within TCGA). Circles: logistic regression; triangles: XGBoost. Lines: cut-off for each model type (midpoint between the two designs; solid, logistic regression; dashed, XGBoost). False alarms: share of negative-control models above the cut-off.

The genes themselves show the difference (Supplementary Table G). The genes that the shortcut models used most were largely pseudogenes and non-coding RNAs, such as *HSPB1P1, MEG3, NEAT1* and the signal recognition particle RNA, together with genes of normal stroma and stress responses. 13 of the 20 most used genes had a larger source effect than cancer effect. A typical source gene was the pseudogene *EIF5AP4* : it was close to zero in GTEx and expressed at a similar level in TCGA tumours and TCGA normals, in six of the seven tissues (Extended Data Fig. 1c). The matched models relied on genes with a known role in cancer, including the tumour suppressor *CDKN2A*, which acts in the Rb and p53 pathways, the stromal genes *MMP11, COL10A1* and *SPP1*, and genes that are often lower in tumours, such as *ADH1B* and *CLEC3B*; only 4 of their 20 most used genes had a larger source effect than cancer effect.

However, reading the explanations this way needs the true source effect of each gene, and therefore matched normals; a practical check must work from the trained model and its training data alone.

### A label-free shortcut score

In the shortcut design, the source effect and the cancer effect of a gene cannot be separated within one tissue, however, they can be separated across tissues. A difference arises from donors, collection, handling and sequencing, so it should look the same in every tissue, whereas cancer changes differ from tissue to tissue (Methods, Eq. 1). In the training data of the shortcut design, *EIF5AP4* differed between tumours and GTEx normals in the same direction and by a similar amount in six of seven tissues (gene score *s* = 2.0), whereas *FGFR4* changed strongly but in opposite directions in different tissues (*s* = 0.1; Fig. 3d). Weighting the gene score by each model’s explanation gives the source-reliance score *R* (Eq. 3), which needs no matched normals, no source labels and no new data.

Recent studies shows, an AUC that looks ‘too good to be true’, separated shortcut-design from matched-design models. The question is whether it can also tell them from models that are near-perfect for honest reasons. As negative controls, we trained logistic regression and XGBoost on 126 easy tasks per model type, telling two tissues apart within one data source, where no source difference exists. The ‘too perfect’ rule flagged 92% (95% CI 86–96%) of the honest logistic regression models and 87% (80–93%) of the XGBoost models (Fig. 3e). It could hardly tell shortcut models from honest ones (AUC 0.58 and 0.61). *R* flagged none of them (0%, 95% CI 0–3%, for both types; P < 0.001 against the AUC rule, exact McNemar test) and separated shortcut models from honest ones perfectly (AUC 1.00; Fig. 3f). Near-perfect accuracy is therefore not evidence of a shortcut, and its absence is not evidence of a sound model; what the model relies on is.

We compared *R* with existing approaches, grouped by what they need (Table 2). The model only uses, as *R* does, separated the two designs and raised false alarms on the negative controls (90% for the AUC rule, 48% for overconfidence) and ranked the graded models less well (*ρ* = 0.64–0.76, against 0.87 for *R*). The share of new samples called cancer ranked the models best (*ρ* = 0.92), however, they needs a pool of unlabelled samples that already contains the failing group, which is the information that is missing in practice. Methods that need labels for the shortcut attribute^19^ cannot be applied, because that attribute is the label itself.

**Table 2:**
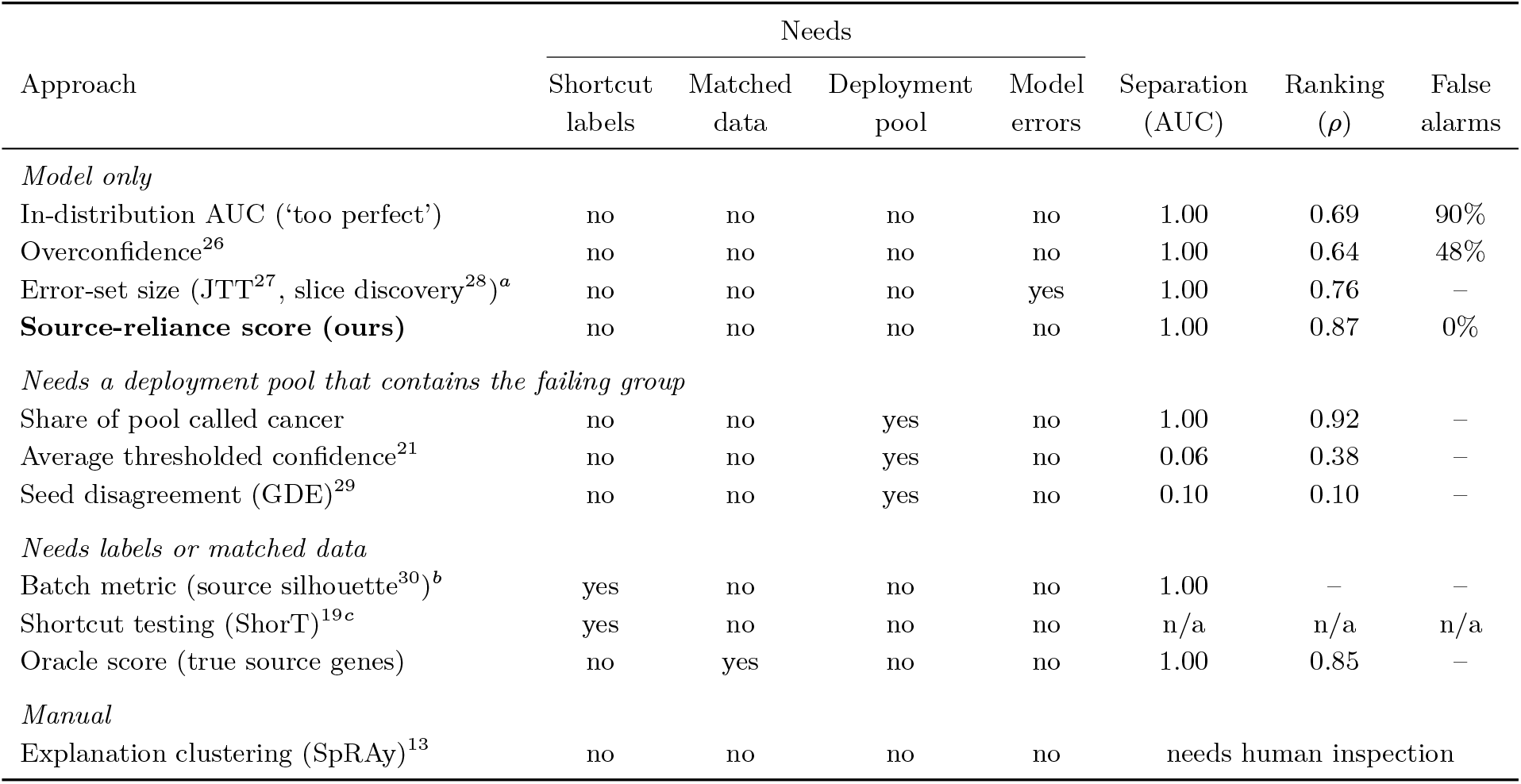
Comparison with existing approaches for detecting a shortcut. Columns 2–5: what each approach needs beyond the trained model and its training data. Separation: AUC for telling shortcut-design models from matched-design models of the same type (mean over model types; *<* 0.5 means the approach ranks the shortcut models as better). Ranking: Spearman correlation with the real failure across the fix series (mean over model types). False alarms: share of negative controls (models trained on easy tasks with no source difference: tissue vs tissue, one data source) that are flagged. ^*a*^Fewer in-distribution errors flags a model; this is the ‘too perfect’ rule in another form, and the error set that JTT and slice discovery work from is empty for shortcut models (0–1 samples). ^*b*^Separation arises only from the genes each design selects; the metric describes the data, not what the model uses. ^*c*^Not applicable: the shortcut attribute (data source) is identical to the label. CPTAC ranking for our score: LR 0.73; XGBoost 0.01 with the cell-division filter, 0.90 without (Table 3).

|  | Needs |  |  |  |  |  |  |
|---|---|---|---|---|---|---|---|
| Approach | Shortcut labels | Matched data | Deployment pool | Model errors | Separation (AUC) | Ranking (*ρ*) | False alarms |
| *Model only* |  |  |  |  |  |  |  |
| In-distribution AUC (‘too perfect’) | no | no | no | no | 1.00 | 0.69 | 90% |
| Overconfidence<sup>26</sup> | no | no | no | no | 1.00 | 0.64 | 48% |
| Error-set size (JTT<sup>27</sup>, slice discovery<sup>28</sup>)<sup>*a*</sup> | no | no | no | yes | 1.00 | 0.76 | – |
| **Source-reliance score (ours)** | no | no | no | no | 1.00 | 0.87 | 0% |
| *Needs a deployment pool that contains the failing group* |  |  |  |  |  |  |  |
| Share of pool called cancer | no | no | yes | no | 1.00 | 0.92 | – |
| Average thresholded confidence<sup>21</sup> | no | no | yes | no | 0.06 | 0.38 | – |
| Seed disagreement (GDE)<sup>29</sup> | no | no | yes | no | 0.10 | 0.10 | – |
| *Needs labels or matched data* |  |  |  |  |  |  |  |
| Batch metric (source silhouette<sup>30</sup>)<sup>*b*</sup> | yes | no | no | no | 1.00 | – | – |
| Shortcut testing (ShorT)<sup>19*c*</sup> | yes | no | no | no | n/a | n/a | n/a |
| Oracle score (true source genes) | no | yes | no | no | 1.00 | 0.85 | – |
| *Manual* |  |  |  |  |  |  |  |
| Explanation clustering (SpRAy)<sup>13</sup> | no | no | no | no | needs human inspection |  |  |

### Validation across models and cohorts

A useful score must follow how much a model relies on the shortcut, not only whether it does. Along the matched-sample ladder, *R* fell with the real failure in all four model types used to develop it (Fig. 4a; Spearman *ρ* = 0.95 for logistic regression, 95% CI 0.81–0.99; 0.94 for XGBoost, 0.75–1.00; 0.86 for TabM, 0.64–0.95; and 0.75 for the MLP, 0.33–0.93; all P < 0.001), and it separated the shortcut and matched designs of every type (AUC 1.00, P < 0.001).

**Figure 4:**
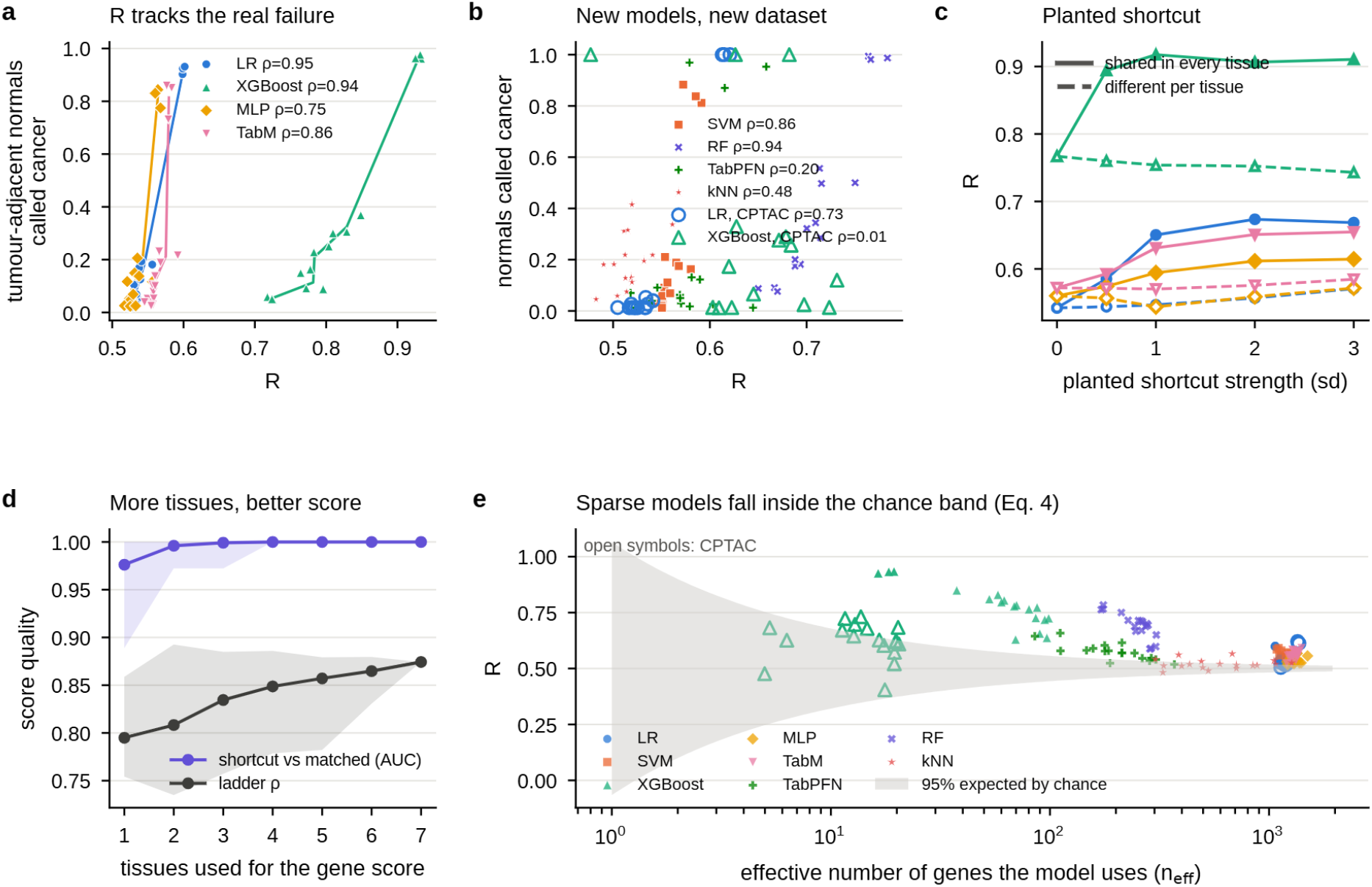
The score tracks real reliance, transfers to new models and data, and fails where its theory predicts. **a**, *R* against the share of tumour-adjacent normals called cancer along the matched-sample ladder (*k* = 0, 5, 10, 20, all; three seeds); lines join means over seeds. *ρ*: Spearman correlation. **b**, The same for held-out model types on TCGA (filled) and for the CPTAC ladder (open). **c**, *R* against the strength of a shortcut planted in the training tumours, either in the same genes in every tissue (solid) or in different genes per tissue (dashed). **d**, Separation of shortcut from matched designs (AUC) and ladder ranking (*ρ*) when the gene score is computed from *m* of the seven tissues; lines: mean over subsets; shading: range. **e**, *R* against the effective number of genes a model uses (*n*_eff_) for every scored model; grey band: 95% range expected by chance (Eq. 4); open symbols: CPTAC.

We then froze *R* and applied it to model types and data that played no part in its development (Fig. 4b). It ranked the ladder of the support vector machine (*ρ* = 0.86, 95% CI 0.57–0.95) and the random forest (*ρ* = 0.94, 0.77–0.99), and that of logistic regression on CPTAC (*ρ* = 0.73, 0.26–0.92). It failed for three: TabPFN (*ρ* = 0.20, *−*0.45 to 0.74), *k*-nearest neighbours (*ρ* = 0.48, *−*0.04 to 0.82), and XGBoost on CPTAC (*ρ* = 0.01, *−*0.62 to 0.59). We examine these failures below.

Two further experiments probed the assumptions behind the score. First, as a positive control, we planted a shortcut of known strength into the training tumours of the matched design; *R* rose with its strength when the same genes were shifted in every tissue, as assumption A1 requires (Fig. 4c): from no plant to the strongest plant, *R* rose by 0.05 (MLP) to 0.14 (XGBoost). When different genes were shifted in each tissue, *R* changed by *−*0.02 to +0.03, although for logistic regression this small rise was still monotone (*ρ* = 0.86). The score therefore responds mainly to shared signals. Second, when we varied the number of strata, the score improved as more tissues were used to compute the gene score Fig. 4d): with one tissue, where the gene score reduces to the ordinary effect size, ranking was weaker (*ρ* = 0.79 on average) and separation in the worst case was 0.89; with four or more tissues, separation was perfect for every subset, and ranking reached 0.87 with all seven.

### Failure modes of the score

The model of the confound states in advance when *R* should fail (Methods, Table 3). The most general prediction comes from the chance level of the score (Eq. 4): when a model spreads its importance over few genes, the band of values expected by chance is wide, and a real reliance on the source cannot be told apart from noise. Figure 4e shows that this is where the failures lie. Logistic regression, the SVM, the MLP and TabM used more than 1,000 genes effectively (*n*_eff_), giving a null standard deviation of at most 0.006, and their scores sat far outside the chance band. XGBoost used 16–99 genes on TCGA and only 5–20 on CPTAC (null s.d. up to 0.13); on TCGA its shortcut models still lay above the band, but on CPTAC its scores fell inside it (*z* = 1.6). TabPFN and *k*-nearest neighbours, explained by single-gene permutation, gave unstable importances; for *k*-nearest neighbours, shuffling the 100 most source-like genes together recovered much of the signal (ranking 0.74, separation 0.78), suggesting that the problem lies in the explanation rather than in the score (an exploratory analysis, designed after the failure was seen).

**Table 3:** Failure modes predicted by the model of the confound, and what we observed. Each row is an assumption of the source-reliance score (Methods), the failure it predicts, the test, and the result. *n*_eff_ : effective number of genes a model uses; null s.d.: standard deviation of the score when importance is unrelated to the gene score (Eq. 4).

| Assumption | Predicted failure | Test | Observed |
|---|---|---|---|
| A1: the source effect is shared across tissues | a tissue-specific shortcut is missed | planted signal, shared vs tissue-specific | ranking *ρ* 0.85 (shared) vs 0.35 (tissue-specific) |
| A2: the cancer effect varies across tissues | shared cancer programmes inflate the score; the filter can remove true source genes | cell-division filter on vs off, all tests | CPTAC XGBoost ranking 0.01 → 0.90 without the filter; smooth models lose 0.05–0.09 |
| Enough tissues to estimate the spread | few tissues weaken the score | gene score from 1–7 tissues (127 subsets) | worst-case separation 0.89 (1 tissue) → 1.00 (≥4 tissues) |
| Many genes carry the importance (large *n*<sub>eff</sub>) | sparse models give noisy scores | *n*<sub>eff</sub> and null s.d. per model | XGBoost *n*<sub>eff</sub> 16–99 on TCGA, 5–20 on CPTAC (null s.d. up to 0.13; CPTAC *z* = 1.6) vs LR, SVM, MLP, TabM *n*<sub>eff</sub> > 1,000 (null s.d. ≤0.006) |

The second prediction concerns shared cancer programmes, which violate assumption A2. Cell division is the clearest example, and the frozen score removes these genes. This helped models that use many genes, but on CPTAC it removed two genes, *TARDBP* and *PSMA1*, that carried 35–55% of the importance of XGBoost and were in fact strong source genes. Without the filter, the CPTAC ranking of XGBoost rose from 0.01 to 0.90; we report the frozen version as the primary result and the unfiltered one as a sensitivity analysis. A sparse model that rests on a few genes is thus sensitive to any error in the gene score, as Eq. 4 predicts. Finally, because explanation methods differ in scale, *R* is comparable within a model type but not across types (Fig. 4a); expressing it as a *z*-score against its chance level partly corrected this.

## Discussion

When two data sources supply the two classes, a classifier can be close to perfect on the usual test and still have learned the wrong thing. In cancer-versus-normal classification from TCGA and GTEx, six model families, from logistic regression to a tabular foundation model, reached an AUC of 1.000 yet called 80–97% of tumour-adjacent normal tissue cancer, and 100% on an independently processed cohort. Moving normal samples towards the TCGA source, and nothing else, was enough to make the models call them cancer. Explanations showed that the models relied on genes that separate the two sources, and a handful of matched samples per tissue removed most of the shortcut. Our main result, however, concerns how such a shortcut can be detected when no matched samples exist: near-perfect accuracy could not tell a shortcut from an honestly easy task, whereas a label-free score computed from the model’s explanations could.

Accuracy is not evidence of what a model has learned. The informal rule that an AUC which looks “too good to be true” should raise suspicion has never been tested. On 126 negative controls per model type, easy tasks in which no source difference exists, the rule flagged 87–92% of models, and it separated shortcut models from honest ones barely better than chance. The reverse error is equally possible: a shortcut model need not look suspicious, as the shortcut models in our study reached AUCs that are typical of published cancer classifiers^7,8^. Whether a model is right for the right reason^13,14^ is a property of what it relies on, not of how well it scores, and has to be measured separately. A test set drawn from the same sources cannot measure it, because it contains the same confound; only samples that break the link between label and source, such as tumour-adjacent normals here, reveal the failure.

Explanations can provide this measurement, used as a number rather than only as a picture. Explanation methods are known to reveal Clever Hans behaviour, but mostly through inspection by eye^13,18,22^, and post-hoc explanations can miss a spurious signal that the user does not already suspect^23^. Our score does not ask the user to recognise a spurious gene. It uses a property of the confound itself: a difference between data sources arises from donors, collection, handling and sequencing, and tends to look the same in every tissue, whereas cancer changes differ between tissues. With several strata, the two effects become separable even though the label and the source coincide within each stratum, and the explanation only has to say how much the model uses each gene. This turns an explanation into a number that can be compared across models, tested against a ground truth and given a chance level (Eq. 4). The score ranked models by their real reliance on the source, separated shortcut from matched designs, raised no false alarms on the negative controls, and transferred, frozen, to two model types and a cohort that played no part in its development. Where it failed, the failures matched limits that the model of the confound states in advance: too few genes for the chance band to be narrow, explanations too noisy to rank genes, or a filter that removed true source genes.

Methods that detect shortcuts usually need information that is missing in this setting (Ta-ble 2). Shortcut testing^19^ needs labels for the suspected attribute; here, that attribute is the label itself. Estimators of performance under shift^20,21^ need new data or targeted retraining, and the best-ranking alternative in our comparison, the share of new samples called cancer, needs a pool that already contains the group on which the model fails, which is exactly what is missing in practice. Batch-effect corrections^10,31^ describe the data rather than what a given model uses, and need matched samples to anchor the correction; our results also show that correcting the data is not a substitute for retraining when the model is a tree ensemble. The source-reliance score is complementary to these methods: it needs only the trained model, its training data and its explanation, and it answers a different question, namely how much this model relies on the source.

The score has clear limits. It rests on two assumptions, and it fails when they fail. The first holds only in part: measured with matched normals, about 60% of the source effect was shared across tissues, against 43% of the cancer effect (Supplementary Note B). A source effect that differs between strata violates the first assumption and is invisible to the score; in the positive control, a tissue-specific shortcut raised the score only slightly. A cancer programme shared across tissues violates the second; we removed cell division as a known case, but the filter can also remove a true source gene, as it did for XGBoost on CPTAC, and other shared programmes may exist. The score needs several strata; with one tissue it reduces to an ordinary effect size, and in our data four or more tissues were needed for perfect separation. It inherits the quality of the explanation: for models that use few genes the chance band is wide, and single-gene permutation importance was too noisy for TabPFN and *k*-nearest neighbours. Scores are comparable within a model type, not across types, because explanation methods differ in scale. Our study uses one domain, bulk transcriptomics, two cohorts and seven tissues; the CPTAC replication covered four tissues and two model types. The negative controls were tissue-identity tasks, and tumour-adjacent normal tissue, our measure of true reliance, is itself not identical to healthy tissue^12^; the causal source test, which used GTEx normals only, addresses this concern but does not remove it. Finally, the score was revised once during development, after the planted-shortcut test on the four development model types (Supplementary Note A); the held-out model types and CPTAC are the independent tests.

In practice, our results suggest four steps for any classifier trained on classes from different sources. First, do not read near-perfect accuracy as evidence that a model has learned the intended signal. Second, when the confounded label is observed in several strata, compute the source-reliance score from the model’s explanation, compare it within the model type and against its chance level, and check that the model uses enough features for the score to be informative. Third, if matched samples can be obtained, even a few per stratum, add them to training rather than using them only to correct the data or to move the decision threshold. Fourth, where possible, test on samples that break the link between label and source; they are the only direct measure of what the model has learned.

Nothing in the argument is specific to cancer or to gene expression. Whenever cases and controls come from different hospitals, scanners, biobanks or assays, and the data span several strata such as organs, sites or cohorts, a model can learn the origin of a sample instead of its biology, and the same separation of shared from stratum-specific effects applies. Testing the score in such settings, including medical imaging^15–17^, and with explanation methods that are stable for sparse and in-context models, are the natural next steps. More broadly, as models are increasingly trained on whatever public data can be combined, a quantitative check of what they rely on should become as routine as reporting how well they score.

## Methods

To rigorously evaluate how data-source confounding affects machine-learning classifiers in molec-ular diagnostics, we designed a systematic framework encompassing multi-cohort data integra-tion, architectural evaluation, causal validation experiments, explanation-based scoring, and strict statistical benchmarking. Below, we detail the datasets, model configurations, causal tests, score formulation, and analytical procedures used throughout this study.

### Data sources and cohorts

We asked whether tumour classifiers trained on data in which cancer status and data source coincide learn cancer, the source, or both. We combined TCGA and GTEx, replicated on CPTAC, and built designs in which the source was either confounded with the label or separated from it.

We used the UCSC Toil recompute of TCGA and GTEx, in which all samples were re-quantified from raw reads with the same pipeline (STAR alignment and RSEM quantification)^5^. Expression was used as log_2_(TPM + 1). We kept the seven tissues that have TCGA tumours, TCGA tumour-adjacent normal tissue and GTEx normal tissue: breast, colon, kidney, liver, lung, prostate and stomach. For breast, only samples from female donors were kept, because GTEx breast includes male donors. TCGA tumours were primary tumours, and each TCGA person contributed at most one tumour and one tumour-adjacent normal sample. Tissues with several TCGA cancer types were pooled by organ (kidney: clear cell, papillary and chromophobe carcinoma; lung: adenocarcinoma and squamous cell carcinoma), as were the two GTEx colon sites (sigmoid and transverse). This gave 6,157 samples from 4,999 people in three groups: 4,540 TCGA tumours, 529 TCGA tumour-adjacent normals (‘TCGA normal’) and 1,088 GTEx normals (counts per tissue and group in Supplementary Table A).

For replication we used bulk RNA-seq from the Clinical Proteomic Tumor Analysis Con-sortium^24^, processed by the Baylor College of Medicine (BCM) pipeline (RSEM, upper-quartile normalised, log_2_(*x* + 1)), as distributed by the cptac data package^32^. We used the cohorts that match our tissues: clear cell renal cell carcinoma (kidney), lung adenocarcinoma and lung squa-mous cell carcinoma (lung), colon adenocarcinoma (colon) and breast cancer (breast). CPTAC tumour-adjacent normal samples served as matched normals, and CPTAC tumours were paired with GTEx normals of the same tissue. Genes were matched by name, and duplicated genes were averaged. Because the two pipelines report expression on different scales, each CPTAC sample was quantile-normalised to the average GTEx distribution. This aligns the overall distributions but does not necessarily remove gene-specific differences between the pipelines.

To state the problem precisely, each sample *i* has an expression profile *x*_*i*_, a label *y*_*i*_ *∈ {*0, 1*}* (1 = tumour), a data source *s*_*i*_ ∈ {TCGA, GTEx} and a tissue *t*_*i*_ ∈ {1, …, *T*}, with *T* = 7. Under the shortcut design, a classifier *f* is trained on TCGA tumours and GTEx normals, so that in the training data *y* = **1**[*s* = TCGA]: the label and the source are the same variable. Under the matched design, *f* is trained on TCGA tumours and TCGA normals, so both classes come from one source. We define the true source reliance of a classifier as the share of matched normals that it calls cancer,

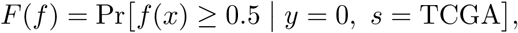

estimated on held-out TCGA normals (the ‘stress test’). A classifier that separates cancer from normal tissue without using the source should have *F* close to 0; one that relies strongly on the source has a higher *F*, up to 1. *F* needs matched normals, which most studies do not have, so we used it only as the reference against which the source-reliance score was judged, never as an input to it. The aim is to estimate reliance on the source from the trained model and its training data alone.

All splits were made by person, so that samples from one person (for example a tumour and its adjacent normal, or several GTEx tissues from one donor) were never split between training and test. For each of five random seeds, 30% of people (1,500 of 4,999; about 1,850 samples) formed the test set. The six main model families were trained on up to five seeds (Fig. 2a,b shows each one); explanations, the score and the matched-sample ladder used seeds 0–2. Within the training set, 15% of people were held out for early stopping. For CPTAC, 30% of patients formed the test set, and GTEx samples kept their usual split. For every model, the 2,000 genes with the highest variance in that model’s training set were selected, and each gene was standardised with the mean and standard deviation of the training set. No test data were used for gene selection or scaling.

In the shortcut design the label and the source cannot be told apart in the training data. We next train a broad range of classifiers on it and ask whether the usual test can reveal what they learned.

### Model architectures and evaluation protocols

We chose six model families that span the main inductive biases for tabular data, from linear to tree ensembles, neural networks and a tabular foundation model. Logistic regression used L2 regularisation (*C* = 0.1) and balanced class weights. The support vector machine used an RBF kernel (*C* = 1, *γ* = ‘scale’), balanced class weights and Platt-scaled probabilities. XGBoost^33^ used up to 1,000 trees (learning rate 0.05, maximum depth 4, row subsampling 0.8, column subsampling 0.3, minimum child weight 2, L2 penalty 1, positive class weight equal to the class ratio), with early stopping after 50 rounds without improvement in validation log-loss. The MLP had two hidden layers (512 and 128 units) with layer normalisation, ReLU and dropout 0.3. TabM^34^ used the default configuration (32 ensemble members, three blocks of 512 units). The MLP and TabM were trained with AdamW (learning rate 10^*−*3^; weight decay 10^*−*4^ and 3 *×* 10^*−*4^), batch size 128, a cosine learning-rate schedule, a class-weighted binary cross-entropy loss, at most 100 epochs and early stopping with a patience of 10 epochs on the validation loss; TabM predictions were averaged over its ensemble members. TabPFN v2.5^25^ was used without training, with the training set as context and balanced class probabilities. Two further model types, a random forest^35^ (300 trees, 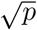 features per split, minimum leaf size 2, balanced class weights) and *k*-nearest neighbours (15 neighbours, distance-weighted), were trained only to test the source-reliance score on model types that played no part in its development; SVM and TabPFN were also held out from development. Models were implemented with scikit-learn^36^, XGBoost, PyTorch^37^ and the tabm and tabpfn packages, in Python 3.13 (NumPy 2.1.3, pandas 2.2.3, SciPy 1.16.3, scikit-learn 1.6.1, XGBoost 3.4.1, PyTorch 2.11, SHAP 0.52). The full environment is listed with the code.

All models were evaluated on the same test set, which contains all three groups. A sample was called ‘cancer’ if its predicted probability was at least 0.5. The usual test is the AUC between tumours and the normals the model was trained against (GTEx normals for the shortcut design). The stress test is the true source reliance *F*. We also report the AUC between tumours and TCGA normals, and the share of GTEx normals and of tumours called cancer.

A large gap between the usual test and the stress test shows that a classifier has learned something other than cancer. To show that this something is the data source, we tested it directly, and we tested what removes it.

### Causal validation and remediation strategies

We first tested the source directly. If shortcut models use it, moving a normal sample towards the TCGA source, and nothing else, should make them call it cancer. For each tissue, we estimated the source shift as the per-gene difference between the means of TCGA normals and GTEx normals in the training set. Each held-out GTEx normal was moved by *β* times this vector, *β ∈ {*0, 0.25, 0.5, 0.75, 1*}* (*β* = 1 moves it to the TCGA-normal average, without extrapolation), and negative values were set to zero. As a control, the same shift was applied with genes randomly permuted within each tissue, which keeps its size but breaks its direction. We also split the shift into a part along a ‘handling’ direction and the remainder; the handling direction was estimated from GTEx training samples alone as the mean difference between the best-and worst-preserved thirds of samples, ranked by RNA integrity number and ischaemic time.

We then asked how many matched samples remove the shortcut. We added *k* TCGA nor-mals per tissue to the shortcut design’s training set, labelled as normal, with *k ∈ {*0, 5, 10, 20*}* or all available training TCGA normals (‘all’). The added samples were drawn at random from training-set TCGA normals with a fixed seed, and the same samples were used in every com-parison. This series also serves as a graded ladder of models whose true reliance falls with *k*, which we use to validate the score.

We compared retraining with three alternatives that use the same *k* samples: (i) keeping the *k* = 0 model and choosing a new decision threshold, either one global threshold or one per tissue, by Youden’s *J* ^38^ between tumours and the added normals; (ii) correcting the data instead of retraining: per tissue, all TCGA samples were shifted by the difference between the mean of GTEx training normals and the mean of the added normals (‘bridge’ correction); estimating the source effect by per-gene linear regression on tumour status and source and removing it, as in removeBatchEffect^39^, gives the same correction, because with three groups this design is saturated; and (iii) both correction and retraining.

Retraining needs matched samples, and most studies that combine sources have none. We therefore asked whether the trained model itself can reveal that it relies on the source, and opened it with explanations.

### Explanations, score formulation and statistical validation

This section follows one chain of reasoning. We first extract from each trained model which genes it uses (explanations), and measure with matched normals what those genes are, as a reference. We then show how source and cancer effects can be separated without matched normals, using the variation across strata, and turn this into a score for each model (equations (1)–(3)) with a known chance level (equation (4)). Finally, we validate the score in five designed experiments, compare it with existing approaches and quantify the uncertainty of every result.

Each model was explained on held-out test samples, and the global importance of a gene was its mean absolute attribution over samples. We used the standard method for each family: weight *×* input for logistic regression, TreeSHAP for XGBoost^40^, and Integrated Gradients^41^ for the MLP and TabM (zero baseline, which is the training mean after standardisation; 32 steps; at most 400 samples). For the SVM, random forest, *k*-nearest neighbours and TabPFN we used permutation importance: the mean absolute change in predicted probability when one gene was shuffled across samples (150 samples; 40 for TabPFN, for speed). For robustness, the main comparison was repeated with permutation importance, XGBoost gain, and, for the MLP and TabM, gradient *×* input and SmoothGrad.

To check what the models rely on, we computed for each gene and tissue Cohen’s *d* (difference in means divided by the pooled standard deviation) on training samples. The true source effect of a gene is the mean over tissues of |*d*| between TCGA normals and GTEx normals; its cancer effect is the mean of |*d*| between TCGA tumours and TCGA normals. A model’s reliance on source genes (or cancer genes) is the importance-weighted mean percentile of the true source effect (or cancer effect) among its genes. These quantities need matched normals and were used only for checking, never by the source-reliance score.

Read this way, explanations reveal the shortcut only when the source genes are already known, which again needs matched normals. The score below removes that need.

The score *R* estimates how strongly a trained model relies on genes that separate the data sources. It uses only what the model was trained on (tumours from one source, normals from another) and the model’s explanation. It needs no matched normals, no labels for the source and no deployment data.

Let *d*_*g*,*t*_ be the standardised difference (Cohen’s *d*) of gene *g* between tumours and GTEx normals in tissue *t*, computed on training samples. In the shortcut design this difference mixes two effects,

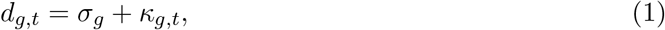

where *σ*_*g*_ is the effect of the data source and *κ*_*g*,*t*_ is the effect of cancer. Within a single tissue the two terms cannot be separated: this is the shortcut, and it is why matched normals are usually thought to be necessary. We make two assumptions. (A1) The source effect is shared: it is the same in every tissue, because it arises from donor type, sample collection, handling and sequencing centre rather than from the tissue. (A2) The cancer effect is heterogeneous: it varies across tissues, with spread *τ*_*g*_ = sd_*t*_(*κ*_*g*,*t*_) *>* 0, and its mean over tissues, 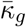, is small compared with *σ*_*g*_ for source genes.

Under (A1) and (A2) the two effects become separable once several tissues are available. Averaging equation (1) over *T* tissues gives 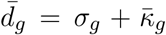, and the spread over tissues gives sd_*t*_(*d*_*g*,*t*_) = *τ*_*g*_, because the shared term does not vary. A gene whose difference is large on average but nearly constant across tissues is therefore source-like; a gene whose difference changes from tissue to tissue is cancer-like. This separation rests on (A1) and (A2); on its own it does not prove that every shared difference is caused by the data source. The tissues play the role of strata, and nothing in the argument is specific to tissues: any design in which the confounded label is observed in several strata (hospitals, scanners, cohorts, assay batches) admits the same separation.

We estimate how source-like each gene is by the signal-to-noise ratio of the shared component,

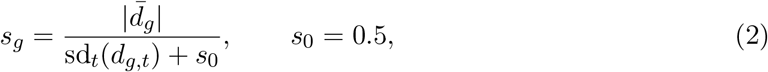

where the constant *s*_0_ keeps the denominator positive and prevents genes with a near-zero spread from dominating, as the fudge factor does in significance analysis of microarrays^42^. It was fixed before any test and frozen. Recomputing *R* with *s*_0_ from 0.1 to 2, a 20-fold range, did not change the separation of shortcut-design from matched-design models (AUC 1.00 for every development model type) and changed the difference between them by at most 0.006 (Spearman *ρ ≥* 0.999 with the values at *s*_0_ = 0.5; Supplementary Table F). The main known violation of (A2) is a cancer programme that is shared across tissues, so that 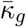 is large and *τ*_*g*_ small; cell division is the clearest such programme. We therefore set *s*_*g*_ = 0 for genes in the MSigDB Hallmark E2F targets, G2/M checkpoint, mitotic spindle and MYC targets gene sets^43^ (the ‘cell-division filter’; 2.2 % of model genes on average). This is a prior on a known violation, not a fitted quantity.

The gene score is then combined with the model’s explanation. Let *w*_*g*_ *≥* 0 be the model’s global importance of gene *g* (negative values set to zero), normalised so that ∑_*g*_ *w*_*g*_ = 1, and let *p*_*g*_ = (rank(*s*_*g*_) *−* 0.5)*/G* be the percentile of the gene score among the model’s *G* genes; ties, which arise mainly from the filtered genes (*s*_*g*_ = 0), receive the average of their ranks. The percentiles therefore lie strictly between 0 and 1, from 1*/*(2*G*) to 1 *−* 1*/*(2*G*). The source-reliance score is the importance-weighted mean percentile,

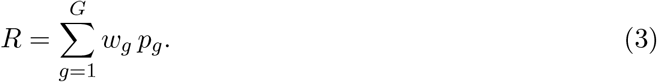

Using percentiles rather than raw scores makes *R* independent of the scale of *s*_*g*_ and robust to a few extreme genes. Because the weights sum to 1, *R* also lies strictly between 0 and 1; it approaches 1 when the model puts all its importance on the most source-like genes.

To know what value of *R* to expect by chance, note that if importance is unrelated to the gene score, the percentiles are a random permutation of 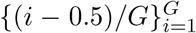 with respect to the weights (ties leave the mean unchanged and reduce the variance slightly), and

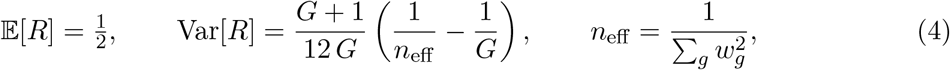

which follows from the variance of a weighted sum under sampling without replacement (checked by simulation). Here *n*_eff_ is the effective number of genes the model uses. Equation (4) gives the band of values expected by chance for each model, and a *z*-score, 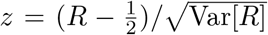.

We use *z* only to judge whether a value of *R* can be told apart from chance, not as a shortcut threshold: a model that relies on cancer genes can also be far from chance.

Equations (1)–(4) state in advance when *R* should fail, and each prediction was tested: (i) a source effect that differs between tissues violates (A1), so a tissue-specific shortcut should be missed (tested with planted tissue-specific signals); (ii) a shared cancer programme outside the filter violates (A2) and inflates the score of cancer genes, while a filter that removes a true source gene deflates it; (iii) with few tissues the spread over tissues is poorly estimated (tested by recomputing the gene score from subsets of tissues); and (iv) a model that uses few genes, or an explanation that is noisy, gives a small or unstable *n*_eff_, so *R* falls inside the chance band of equation (4).

For evaluation only, we computed an oracle score by replacing *s*_*g*_ with the true source effect, which requires matched normals. The oracle gives the best that any explanation-based score could achieve with these explanations.

An earlier version of the score, the Spearman correlation between importance and gene score, failed when a model put all its importance on a few genes (Supplementary Note A). We replaced it with *R* after the planted-shortcut test and froze *R*, including the filter and *s*_0_, before scoring TabPFN, the random forest, *k*-nearest neighbours and CPTAC. All tests were also rescored without the cell-division filter as a sensitivity analysis; the frozen version is the primary result. A score of source reliance can only be validated on models whose reliance is known. We therefore validated *R* in five experiments in which the ground truth is set by the design rather than estimated. Each tests one property that a valid score must have: a monotonic response to graded reliance, resistance to false positives, transfer to unseen models and cohorts, recov-ery of planted signals, and sensitivity to the number of strata. In every experiment *R* was compared only within a model type, because explanation methods differ in scale between types (equation (4)).

The first experiment tested a monotonic response to graded reliance. The matched-sample ladder (*k* = 0, 5, 10, 20 and all; 3 seeds) gives models of the same type whose true reliance *F* falls step by step, so a valid score should fall with it. We measured the Spearman correlation between *R* and *F* within each model type, and the AUC of *R* for separating shortcut-design from matched-design models of the same type.

The second experiment tested resistance to false positives with negative controls, to tell a shortcut from a task that is simply easy. We trained logistic regression and XGBoost on tasks with near-perfect accuracy but no source difference: telling two tissues apart using only GTEx normals, or only TCGA tumours (21 tissue pairs, 2 sources, 3 seeds; 126 models per model type). By construction these models do not rely on the source, so every flag is a false alarm. The shortcut and matched designs were retrained with the same code, and a model was flagged when its value passed the midpoint between the matched-design and shortcut-design means of its model type. The same rule was applied to the in-distribution AUC, the informal ‘too perfect’ check.

The third experiment tested transfer to unseen models and cohorts. After freezing, *R* was applied unchanged to four model types that played no part in its development (SVM, random forest, *k*-nearest neighbours and TabPFN) on the TCGA ladder, and to logistic regression and XGBoost on the ladder of an independent cohort, CPTAC, with the gene score recomputed from CPTAC training data.

The fourth experiment tested recovery of planted signals, using positive controls: shortcuts of known strength. Starting from the matched design, we added a synthetic signal of strength *δ ∈ {*0, 0.5, 1, 2, 3*}* standard deviations to 40 genes in training tumours only. In one arm the genes were the same in every tissue (a shared shortcut, which satisfies A1); in the other they differed between tissues (a tissue-specific shortcut, which violates A1). The gene score was recomputed from the planted training data. The design predicts that *R* rises with *δ* in the shared arm and changes little in the tissue-specific arm; we measured the rise in *R* and its Spearman correlation with *δ*. Because this experiment was also used during development, it is not independent of the final form of the score.

The fifth experiment tested sensitivity to the number of strata. The gene score uses the variation across strata, so its quality should depend on how many are available. We recomputed it from every subset of *m* of the seven tissues (*m* = 1, …, 7; 127 subsets; for *m* = 1 it reduces to |*d*|), rescored the saved explanations and measured separation and ranking as a function of *m*. As robustness checks, we repeated the designs with 2,000 random expressed genes or the 2,000 most highly expressed genes (mean log_2_(TPM + 1) *≥* 1; Supplementary Table B). To understand the failure for *k*-nearest neighbours, we compared single-gene importance with a group permutation in which the 100 genes with the highest gene score were shuffled together, relative to the mean of five random blocks of 100 genes (300 samples); this analysis was designed after the failure and is exploratory. To check assumption A1 directly, we measured the source effect with the matched normals (TCGA normal vs GTEx normal, Cohen’s *d* per tissue, seed-0 training set, expressed genes) and compared its agreement between tissues with that of the cancer effect (Supplementary Note B, Supplementary Table E); this check uses matched normals and plays no part in the score.

Unlike most methods for detecting shortcuts, *R* needs no labels for the shortcut attribute, no matched samples and no deployment data; it separates the source from the label by using the variation across strata. We compared *R* with approaches grouped by what they need beyond the trained model and its training data (Table 2). Model-only approaches use the same information as *R*: the in-distribution AUC (‘too perfect’), the mean confidence |2*p −* 1| on in-distribution test samples^26^, and the number of in-distribution errors, which is the set that JTT^27^ and slice discovery^28^ work from. Deployment-pool approaches need a set of new unlabelled samples that contains the failing group: the share of the pool called cancer, average thresholded confidence^21^, and the disagreement between two seeds^29^. Approaches that need labels or matched data: the silhouette of the data source in the model’s input genes^30^, a batch-effect metric that needs source labels; shortcut testing (ShorT)^19^, which needs labels for the shortcut attribute and is not applicable here because that attribute is the label itself; and the oracle score. Explanation clustering (SpRAy)^13^ needs human inspection and was not scored. Each approach was scored with the same separation, ranking and false-alarm criteria as *R*, and with its sign chosen so that higher values mean more suspicion.

Finally, we quantified how certain each of these results is.

Results are means over seeds, with individual seeds shown in the figures. Because different seeds re-use many of the same test samples, seeds were not treated as independent replicates. Rates (for example, the share of normals called cancer) are reported with Wilson 95% confidence intervals using the number of test samples of a single seed, which is conservative. Shortcut and matched designs, the source shift and its random control, and retraining and correction were compared with one-sided or two-sided Fisher’s exact tests on the same counts. AUCs are reported with 95% confidence intervals from 2,000 bootstrap resamples of test samples, stratified by class and averaged over seeds. Spearman correlations between *R* and true reliance are reported with 95% bootstrap confidence intervals over trained models and one-sided permutation *P*-values (10,000 permutations). Separation of shortcut-design from matched-design models was tested by permuting design labels within each model type. False-alarm rates are reported with exact Clopper–Pearson intervals, and the AUC rule and *R* were compared on the same models with an exact McNemar test. All values are listed in Supplementary Table C. All tests were two-sided unless stated otherwise.

## Data availability

TCGA and GTEx data are available from the UCSC Xena Toil recompute (https://xenabrowser.net). CPTAC data are available through the cptac package (Zenodo record 8394329). The processed data and all saved results needed to reproduce the figures and tables (sample table, person-level splits, expression matrix, model results, test scores and explanations) are available on Zenodo (doi:10.5281/zenodo.23474750). Source Data for Figs. 1–4 and Extended Data Figs. 1 and 2 are provided with this paper.

## Code availability

The notebook reproduce_paper.ipynb, which regenerates every figure, table and number in the paper and its Supplementary Information from the saved results on CPU, and the notebook used to train all models are available on Zenodo (doi:10.5281/zenodo.23474750).

## Extended Data

**Extended Data Fig. 1.**
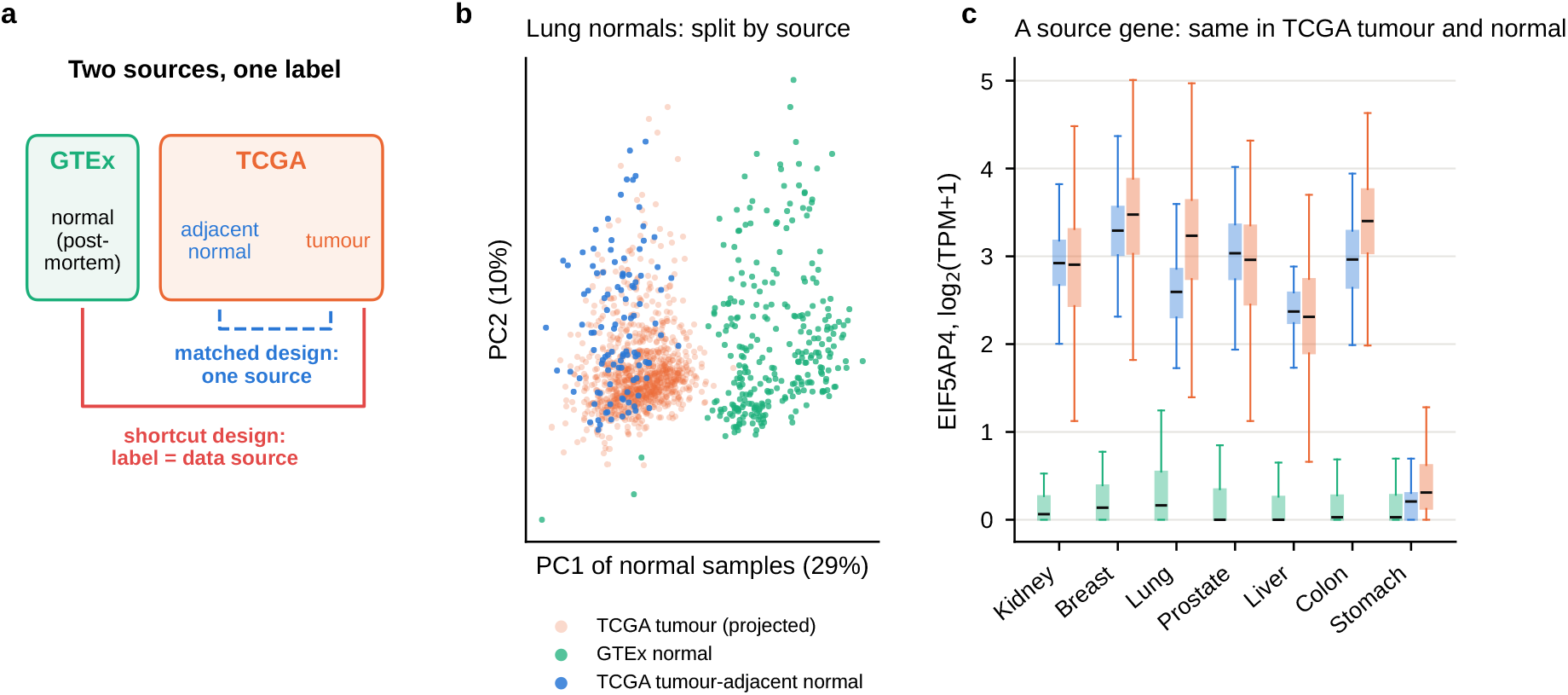
Two data sources, one label. **a**, In the shortcut design (red), tumours from TCGA are compared with normals from GTEx, so the label and the data source coincide. In the matched design (blue), both classes come from TCGA. **b**, Principal components of lung normal tissue only, fitted on GTEx and TCGA normals; TCGA tumours are projected. Normal tissue from the two sources separates along the first component, and TCGA normals lie with TCGA tumours. **c**, Expression of *EIF5AP4* in each tissue and group. In six of the seven tissues the gene is close to zero in GTEx and expressed at a similar level in TCGA tumours and TCGA normals: a source gene, not a cancer gene. In stomach it is low in both TCGA groups.

**Extended Data Fig. 2.**
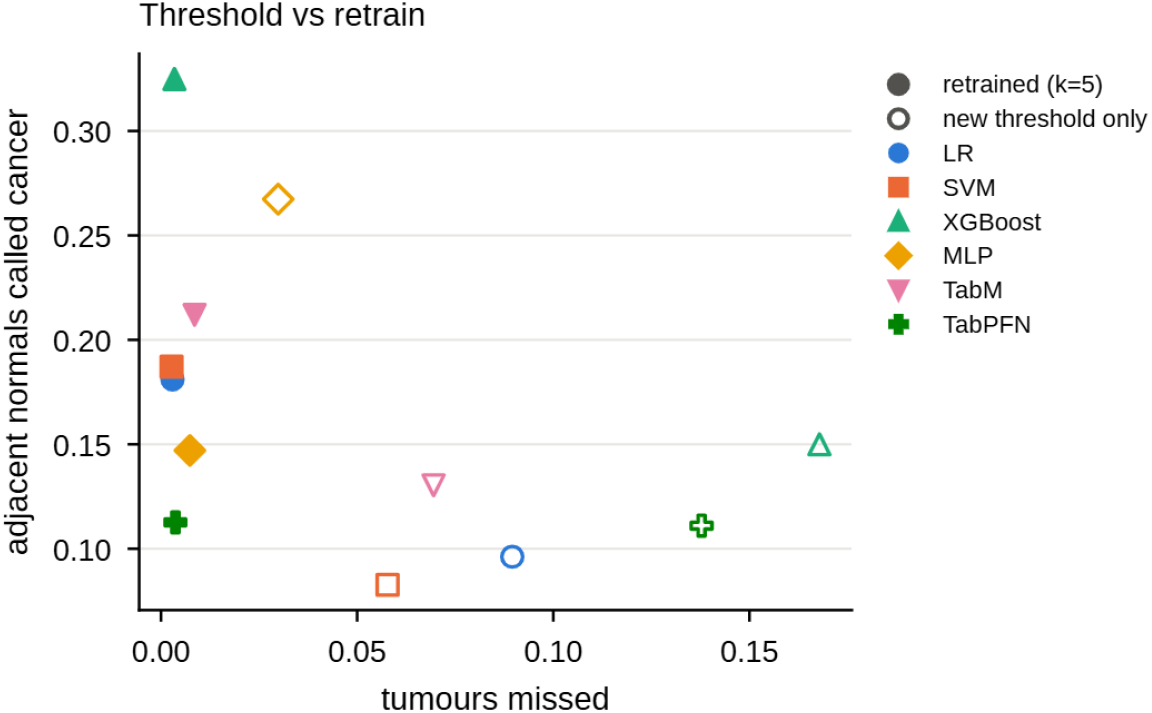
A new threshold trades missed tumours for fewer false calls; retrain-ing does not. Share of tumours missed against share of tumour-adjacent normals called cancer, using the same five matched normals per tissue either to choose a new global decision threshold for the *k* = 0 model (open) or to retrain the model (filled). Colours and shapes: model families as in Fig. 2.

## References

1. Weinstein, J. N. et al. The Cancer Genome Atlas Pan-Cancer analysis project. Nature Genetics 45, 1113–1120 (2013).

2. GTEx Consortium, Lonsdale, J., et al. The Genotype-Tissue Expression (GTEx) project. Nature Genetics 45, 580–585 (2013).

3. Hanahan, D. Hallmarks of cancer: new dimensions. Cancer Discovery 12, 31–46 (2022).

4. Kandoth, C. et al. Mutational landscape and significance across 12 major cancer types. Nature 502, 333–339 (2013).

5. Vivian, J. et al. Toil enables reproducible, open source, big biomedical data analyses. Nature Biotechnology 35, 314–316 (2017).

6. Wang, Q. et al. Unifying cancer and normal RNA sequencing data from different sources. Scientific Data 5, 180061 (2018).

7. Ahn, T., Goo, T., Lee, C.-h., Kim, S., Han, K., Park, S. & Park, T. Deep learning-based identification of cancer or normal tissue using gene expression data. In 2018 IEEE International Conference on Bioinformatics and Biomedicine (BIBM) 1748–1752 (2018).

8. Hong, J., Hachem, L. D. & Fehlings, M. G. A deep learning model to classify neoplastic state and tissue origin from transcriptomic data. Scientific Reports 12 (2022).

9. Quinn, T. P., Nguyen, T., Lee, S. C. & Venkatesh, S. Cancer as a tissue anomaly: classifying tumor transcriptomes based only on healthy data. Frontiers in Genetics 10, 599 (2019).

10. Leek, J. T. et al. Tackling the widespread and critical impact of batch effects in high-throughput data. Nature Reviews Genetics 11, 733–739 (2010).

11. Soneson, C., Gerster, S. & Delorenzi, M. Batch effect confounding leads to strong bias in performance estimates obtained by cross-validation. PLoS ONE 9, e100335 (2014).

12. Aran, D. et al. Comprehensive analysis of normal adjacent to tumor transcriptomes. Nature Communications 8, 1077 (2017).

13. Lapuschkin, S. et al. Unmasking Clever Hans predictors and assessing what machines really learn. Nature Communications 10, 1096 (2019).

14. Geirhos, R. et al. Shortcut learning in deep neural networks. Nature Machine Intelligence 2, 665–673 (2020).

15. Zech, J. R. et al. Variable generalization performance of a deep learning model to detect pneumonia in chest radiographs: a cross-sectional study. PLoS Medicine 15, e1002683 (2018).

16. DeGrave, A. J., Janizek, J. D. & Lee, S.-I. AI for radiographic COVID-19 detection selects shortcuts over signal. Nature Machine Intelligence 3, 610–619 (2021).

17. Howard, F. M. et al. The impact of site-specific digital histology signatures on deep learning model accuracy and bias. Nature Communications 12, 4423 (2021).

18. Kauffmann, J., Dippel, J., Ruff, L., Samek, W., Müller, K.-R. & Montavon, G. Explainable AI reveals Clever Hans effects in unsupervised learning models. Nature Machine Intelligence 7, 412–422 (2025).

19. Brown, A. et al. Detecting shortcut learning for fair medical AI using shortcut testing. Nature Communications 14, 4314 (2023).

20. Ong Ly, C. et al. Shortcut learning in medical AI hinders generalization: method for estimating AI model generalization without external data. npj Digital Medicine 7, 124 (2024).

21. Garg, S., Balakrishnan, S., Lipton, Z. C., Neyshabur, B. & Sedghi, H. Leveraging unlabeled data to predict out-of-distribution performance. In International Conference on Learning Representations (ICLR) (2022).

22. Anders, C. J. et al. Finding and removing Clever Hans: using explanation methods to debug and improve deep models. Information Fusion 77, 261–295 (2022).

23. Adebayo, J., Muelly, M., Abelson, H. & Kim, B. Post hoc explanations may be ineffective for detecting unknown spurious correlation. In International Conference on Learning Representations (ICLR) (2022).

24. Edwards, N. J. et al. The CPTAC Data Portal: a resource for cancer proteomics research. Journal of Proteome Research 14, 2707–2713 (2015).

25. Hollmann, N. et al. Accurate predictions on small data with a tabular foundation model. Nature 637, 319–326 (2025).

26. Guillory, D., Shankar, V., Ebrahimi, S., Darrell, T. & Schmidt, L. Predicting with confidence on unseen distributions. In Proceedings of the IEEE/CVF International Conference on Computer Vision (ICCV) 1114–1124 (2021).

27. Liu, E. Z., Haghgoo, B., Chen, A. S., Raghunathan, A., Koh, P. W., Sagawa, S., Liang, P. & Finn, C. Just train twice: improving group robustness without training group information. In Proceedings of the 38th International Conference on Machine Learning (ICML) 6781–6792 (2021).

28. Eyuboglu, S. et al. Domino: discovering systematic errors with cross-modal embeddings. In International Conference on Learning Representations (ICLR) (2022).

29. Jiang, Y., Nagarajan, V., Baek, C. & Kolter, J. Z. Assessing generalization of SGD via disagreement. In International Conference on Learning Representations (ICLR) (2022).

30. Rousseeuw, P. J. Silhouettes: a graphical aid to the interpretation and validation of cluster analysis. Journal of Computational and Applied Mathematics 20, 53–65 (1987).

31. Molania, R. et al. Removing unwanted variation from large-scale RNA sequencing data with PRPS. Nature Biotechnology 41, 82–95 (2023).

32. Lindgren, C. M. et al. Simplified and unified access to cancer proteogenomic data. Journal of Proteome Research 20, 1902–1910 (2021).

33. Chen, T. & Guestrin, C. XGBoost: a scalable tree boosting system. In Proceedings of the 22nd ACM SIGKDD International Conference on Knowledge Discovery and Data Mining 785–794 (2016).

34. Gorishniy, Y., Kotelnikov, A. & Babenko, A. TabM: advancing tabular deep learning with parameter-efficient ensembling. In International Conference on Learning Representations (ICLR) (2025).

35. Breiman, L. Random forests. Machine Learning 45, 5–32 (2001).

36. Pedregosa, F. et al. Scikit-learn: machine learning in Python. Journal of Machine Learning Research 12, 2825–2830 (2011).

37. Paszke, A. et al. PyTorch: an imperative style, high-performance deep learning library. In Advances in Neural Information Processing Systems 32 8024–8035 (2019).

38. Youden, W. J. Index for rating diagnostic tests. Cancer 3, 32–35 (1950).

39. Ritchie, M. E. et al. limma powers differential expression analyses for RNA-sequencing and microarray studies. Nucleic Acids Research 43, e47 (2015).

40. Lundberg, S. M. et al. From local explanations to global understanding with explainable AI for trees. Nature Machine Intelligence 2, 56–67 (2020).

41. Sundararajan, M., Taly, A. & Yan, Q. Axiomatic attribution for deep networks. In Proceedings of the 34th International Conference on Machine Learning (ICML) 3319–3328 (2017).

42. Tusher, V. G., Tibshirani, R. & Chu, G. Significance analysis of microarrays applied to the ionizing radiation response. Proceedings of the National Academy of Sciences of the USA 98, 5116–5121 (2001).

43. Liberzon, A. et al. The Molecular Signatures Database (MSigDB) hallmark gene set collection. Cell Systems 1, 417–425 (2015).

